# Modelling the emergence of spiral colony morphology in the yeast *Magnusiomyces magnusii*

**DOI:** 10.1101/2025.09.07.674689

**Authors:** Kai Li, Andrew J. Black, Tea Knežević, Jennifer M. Gardner, Jin Zhang, Vladimir Jiranek, J. Edward F. Green, Benjamin J. Binder, Alexander K. Y. Tam

## Abstract

Yeast species have several adaptations that enable them to survive in harsh environments. These adaptations include biofilm formation, where the secretion of extracellular polymeric substances can protect the cells from a hostile environment, or, under nutrient-limited conditions, pseudohyphal or hyphal growth, where the colony can send out long ‘tendrils’ to explore the environment and seek nutrients. Recently, we observed a spiral colony morphology emerge in an isolate of the hyphae-forming yeast *Magnusiomyces magnusii* grown under laboratory conditions. We use an off-lattice agent-based model (ABM) that simulates colony development to investigate the hypothesis that bias in the angle between successive hyphal segments causes the spiral morphology. The model involves biologically-motivated rules of hyphal extension, with key model parameters including the colony size at the onset of hyphal filaments, and the angle between the penultimate and the apical segments. Using one example of an experimentally-grown colony, we use a sequential neural likelihood method to perform likelihood-free Bayesian inference to infer the model parameters. Our results indicate a mean angle between hyphal segments of 2.3^◦^ [1.1^◦^, 3.6^◦^] (95% credible interval). To confirm the model’s applicability to colony growth, we use biologically-feasible parameter values to yield morphologies observed in *M. magnusii* experiments.

## 1 Introduction and background

Yeasts are unicellular eukaryotic organisms that have diverse impacts on human life. A major negative impact is their ability to colonise implanted medical devices such as catheters, prostheses, and stents [35, 42]. Yeasts such as *Candida albicans* and *Candida auris* are opportunistic pathogens responsible for up to 44% of severe hospital-acquired infections [59], which have mortality rates of between 15–35% [53, 2]. Yeasts are infectious because they can invade surrounding tissue [20], thanks, at least in part, to their ability to form hyphae [12] or pseudohyphae [28, 9]. Therefore, some scientific work on yeasts has focused on understanding cellular behaviours that give rise to macroscopic patterns in pseudohyphal [38] or hyphal [43, 44] colonies. Much of this work involves the species *Saccharomyces cerevisiae*, which was the first eukaryotic organism to have its genome fully sequenced [29]. Since then, this yeast has become an even more common model organism in cell biology research [11, 13], aiding the understanding of pathogenic yeasts [8] and the behaviour of eukaryotic cells more generally [13].

*S. cerevisiae* also has uses in the production of food (as reflected by its common name of ‘baker’s yeast’), medicines [1], and biofuels [40]. Another common use of yeast is in brewing alcoholic beverages, where optimisation of yeast strains is an ongoing area of innovation in wine production [26, 64]. Strain optimisation increasingly involves bioprospecting, the process of uncovering new strains in natural habitats, as winemakers seek improved quality and fermentation efficiency [33]. This continual discovery of new strains motivates ongoing scientific research into understanding differences in growth between yeast species and strains in different experimental conditions.

The diversity of yeasts’ impacts on life is mirrored by their diversity in modes of growth. Many different growth modes can be reproduced in laboratory experiments, as illustrated in Figure 1. These morphologies are mediated by factors including the yeast species, genetic make up [58], adhesins [73], nutrient availability [71, 68, 61, 28], quorum sensing [32, 49] and the mechanical properties of the medium on which they are grown [54, 56]. On soft substrates, yeasts can form colony biofilms that can grow over a 90 mm Petri dish in 1–2 weeks [4, 6, 57, 58, 61]. This form of growth is illustrated in Figures 1a–1d. The distributions of viable and dead cells within these colonies can vary spatially [55, 70], and understanding the roles of cell death in yeast colonies is an important subject of research [16, 50, 30, 69]. Yeasts can enter a stationary phase that enables them to survive [46] harsh environments. In seemingly opposite behaviour, yeasts can also undergo regulated cell death [17], an altruistic process that releases nutrients for future consumption [48]. One method for characterising cell death with colonies is to stain them with Phloxine B [45] dye. As illustrated in Figures 1a and 1b, this staining creates darker pink regions indicating elevated cell death. In Figure 1a, cell death occurs near the centre of the colony. This death is likely to be accidental cell death due to lack of nutrients, similar to a necrotic core. Conversely, in Figure 1b cell death is localised in a ring trailing the leading edge. This death may be due to regulated cell death [48], which releases nutrients near the leading edge to promote further proliferation. These different forms of cell death highlight yeast’s diversity of growth.

**Figure 1:**
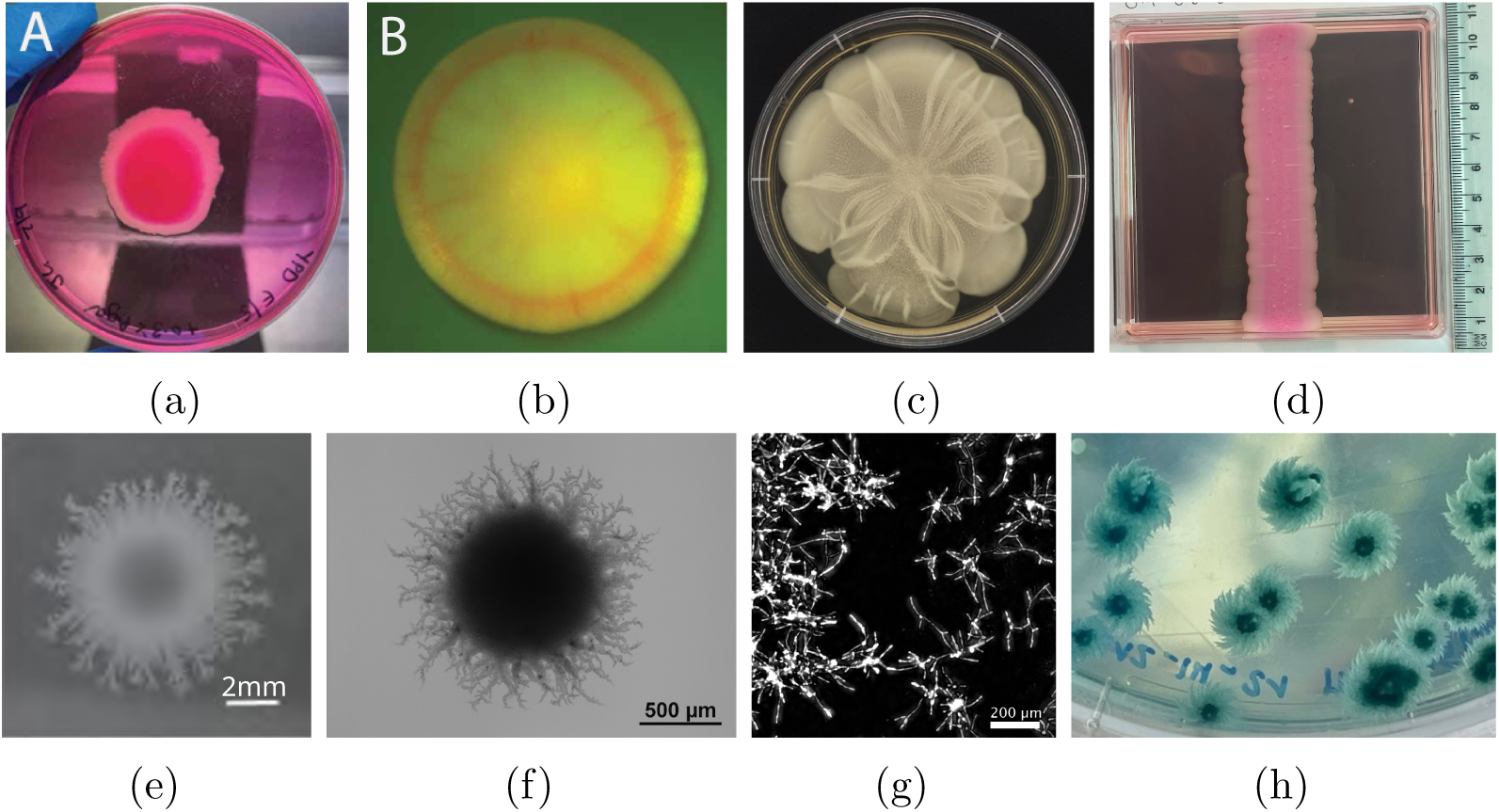
Comparison of yeast colony patterns in different experimental environments. (a) *M. magnusii* colony biofilm after seven days of growth. Dark regions indicate elevated cell death occurring in a central region resembling a necrotic core, which probably occurs due to accidental cell death (ACD) [48]. (b) *S. cerevisiae* colony biofilm after five days of growth. The red ring indicates elevated cell death, which is probably due to regulated cell death (RCD) [48]. (c) *S. cerevisiae* colony biofilm grown on nutrient-rich soft (0.3%) agar [57, 58, 61]. (d) Rectangular *S. cerevisiae* colony biofilm on (0.6%) YPD media. Darker regions indicate elevated cell death compared to lighter regions [62] (e) *S. cerevisiae* colony grown on YND after 58 days of growth [67]. (f) Filamentous yeast of the *S. cerevisiae* strain AWRI 796, with 50 µm nutrients [9]. (g) Intermediate stages of hyphal colony formation in *Candida albicans* [39] (h) Hyphal *M. magnusii* colonies grown on a Petri dish with high colony density.

Colony biofilm growth is also influenced by nutrient availability [61, 71]. In low-nutrient environments, yeasts form very different morphologies to colony biofilms by producing smaller colonies with non-uniform spatial structure (Figures 1e and 1f). Interestingly, Tronnolone et al. [68] found that yeasts, which are non-motile, also cannot respond actively to nutrient gradients. This provided evidence that the non-uniform patterns in Figures 1e and 1f form due to pseudohyphal growth [28, 9, 67], rather than directed growth in response to environmental gradients. Pseudohyphal growth involves a switch between unipolar and bipolar budding and the creation of elongated chains of unipolar cells [28], allowing the colony to extend radially to obtain nutrients, and to invade agar surfaces. Similar to moulds, yeasts can also form hyphal colonies, where the yeast grows in elongated tubes consisting of multiple cells. Figure 1g depicts the early stages of hyphal colony development. Hyphal growth is also affected by many factors, for example carbon dioxide levels [52]. These different colony morphologies illustrate that diversity of outcome is a key characteristic of yeast growth. Given that new strains are being continuously discovered by bioprospecting, there is an incentive for developing robust mathematical modelling and parameter inference methods to understand these new strains. We outline such a procedure in this manuscript.

In search of improved wine strains, we bioprospected a new isolate of *M. magnusii* from the sap of a Tasmanian cider gum tree, *Eucalyptus gunnii*. Under laboratory conditions, this yeast formed a novel highly ordered spiral morphology (Figure 1h). Spiral patterns have been observed during hyphal growth of other fungi [21, 41, 65, 7]. However, the *M. magnusii* morphology is distinct from these other fungi, and features thicker, more densely-packed spirals that form a galaxy-like pattern. We focus on modelling this distinct new morphology. The spiralling morphology in *M. magnusii* does not occur in all growth environments (data not shown). Spiral formation occurred on plates with high colony density, but not on plates with low colony density. Since colonies on high-density plates face more competition for nutrients from the other colonies, spiral formation appears to represent another adaptation that a yeast has developed in response to a harsh environment.

Like *S. cerevisiae*, *M. magnusii* colonies create filaments that extend radially from a circular Eden-like [24] region. Unlike *S. cerevisiae*, where the filaments consist of elongated yeast cells, the main branch of the *M. magnusii* filament consists of stiff rod-like hyphae [22] that are 7–12 µm wide. Penicillate secondary branches emerge at acute angles from the main branch, forming a bushy or fan-like arrangement. These thinner secondary branches break apart into arthroconidia, which are rectangular spore-like units 4–7 µm wide and 10–18 µm long (Figure 2). Non-uniform spatial patterns also emerge in hyphal *C. albicans* colonies [8, 19, 47], but they do not exhibit the striking spiral patterns of *M. magnusii*. The microscale mechanisms that underpin these spiral colonies of *M. magnusii* remain to be determined.

**Figure 2:**
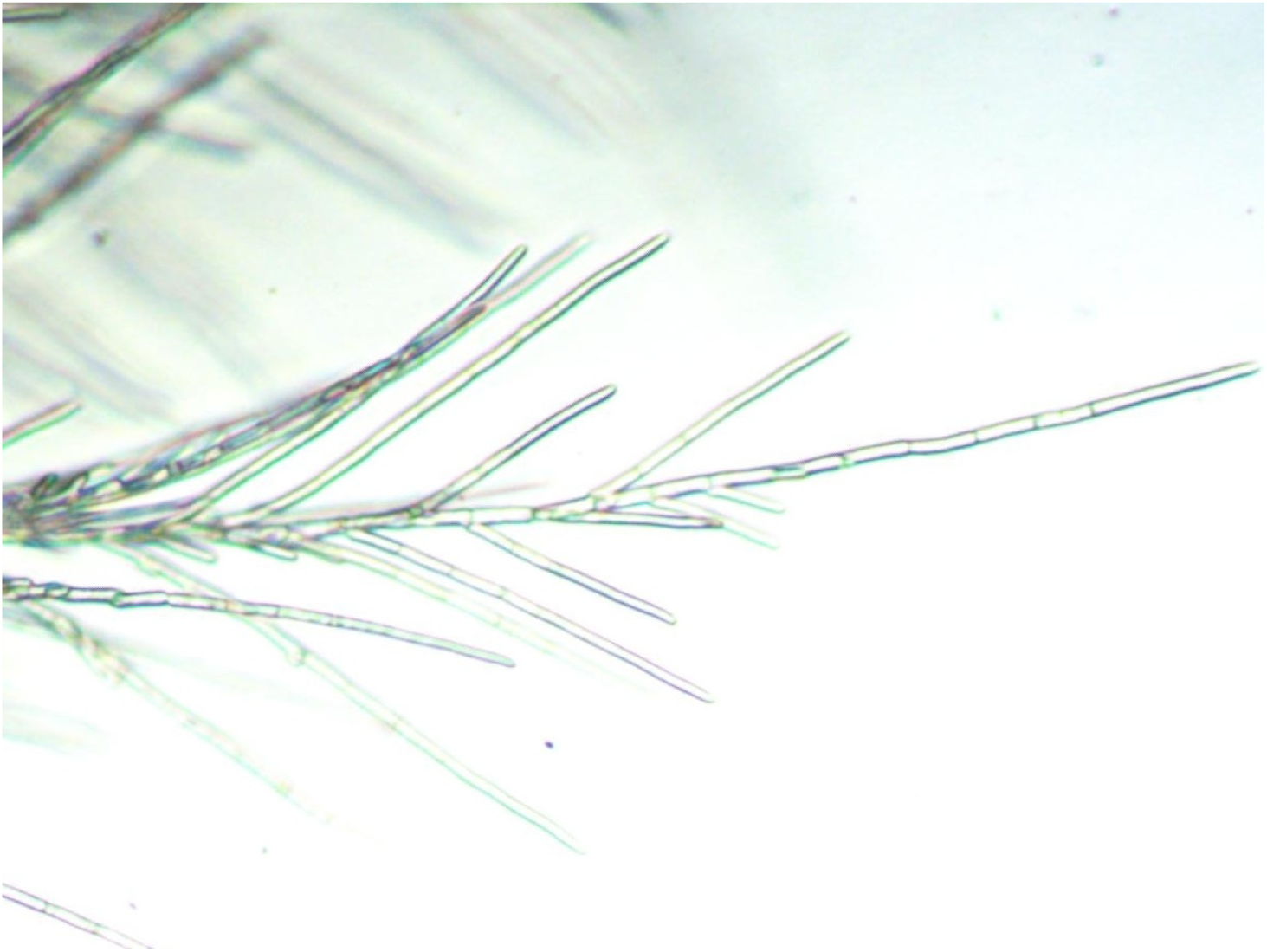
Zoomed view of the the filamentous region of a hyphal *M. magnusii* colony from our experiments.

Depending on the type of growth considered, mathematical models of yeast colonies have involved agent-based [38, 5], reaction–diffusion [18, 61, 36], and continuum mechanical [60, 63] approaches. Owing to the small size of individual *M. magnusii* colonies, we use an off-lattice agent-based model [67, 25, 14, 34, 15, 72] to simulate the spiral morphology. Our objective is to understand how cellular behaviour gives rise to colony-scale patterns. Agents in the model represent rod-like segments of fixed size. These segments could represent hyphae, segments of a rod-like hyphal filament, or arthroconidia in a secondary branch. In the model, regular hyphae and arthroconidia can extend from sites located on the sides of the cell, whereas rod-like (filament-forming) hyphae extend end-to-end only, with a prescribed angle between segments. To compare the model with the experiment, we use image processing and Bayesian inference using neural likelihood estimation [51, 3, 23]. The shape of the hyphal colony components is mainly governed by the angle between successive hyphal segments. Using realistic parameter estimates, the model can also capture three other distinct colony morphologies formed by *M. magnusii* grown under different conditions.

## 2 Yeast growth experiments

An in-house isolate of *Magnusiomyces magnusii* was used for these experiments. Yeast was cultured in liquid Yeast Peptone Dextrose medium (YPD; 10 g L^−1^ yeast extract, 20 g L^−1^ peptone, 20 g L^−1^ glucose) with 2% agar where solid medium was required. 5 mL yeast cultures were grown for 48 hours in liquid YPD prior to inoculation onto 90 mm round agar plates where a 50 µL aliquot diluted to 4 × 10^2^ cells mL^−1^ (low cell density) or 4 × 10^3^ cells mL^−1^ (high cell density) with Phosphate Buffered Saline was spread plated. Agar plates contained either Chemically Defined Sap Medium (CDS; 10 g L^−1^ yeast extract, 20 g L^−1^ peptone, 15 g L^−1^ glucose, 12 g L^−1^ fructose, 9 g L^−1^ maltose, 2 g L^−1^ glycerol, 1% (v/v) ethanol, 1 g L^−1^ acetic acid, 1 g L^−1^ pyruvic acid, 2 g L^−1^ gluconic acid, 0.1 g L^−1^ succinic acid) or Yeast Nitrogen Base (YNB; Becton Dickinson; Cat No. 233520), prepared according to the manufacturer’s instructions and with the addition of 2 g L^−1^ D-glucose and 5 g L^−1^ ammonium sulfate. Plates were sealed with Parafilm and incubated at 25 ^◦^C for 5-10 days. Macroscopic plate images were captured with an Apple iPhone 12 Pro and microscopic imaging with a Leica M-Z FL111 microscope with a Nikon DS-5MC Microscope Camera and Nikon NIS element software or a Carl Zeiss Axiophot Pol Photomicroscope with a Toupcam UCMOS Microscope Camera and ToupView software.

## 3 Mathematical modelling and parameter inference methods

We aim to develop procedures for mathematical modelling and parameter inference that can apply to a variety of yeast colony morphologies. This section describes off-lattice agent-based modelling and a parameter inference method using sequential neural likelihood estimation. We illustrate these methods on a *M. magnusii* spiral colony. The biological insight gained includes quantification of the angle between segments in spiral-forming colonies, and understanding of how model parameters, and hence cellular behaviour, differs across experimental conditions.

### 3.1 Off-lattice agent-based model

Li et al. [38] developed a two-dimensional off-lattice agent-based model for filamentous *S. cerevisiae* colonies, where the filaments consisted of pseudohyphal cells. We adapt this prior model to *M. magnusii* colonies, which are formed from hyphae. Each hypha consists of many individual cells, forming long and tubular structures much longer than an individual cell, but of similar width. Consequently, hyphae do not proliferate by cellular budding, but still grow and fragment as the colony consumes nutrients. To model growth by hyphal extension, we assume that the colony consists of many rod-like agents as per Figures 2 and 3, and we assume that each agent has the same size. All agents are ellipses with major axis length 22.5 µm and minor axis length 7.5 µm, giving an aspect ratio of 3 [22]. In practice, we represent these ellipses as dodecagons in 2D. These agents can be either regular hyphae, which can produce new hyphal segments at four sites located on their sides at angles of ±7*π/*16 radians from the distal poles of the mother hyphae, or filament-forming hyphae, which can only produce a new hyphal segment at one site located at the distal end. To capture the anticlockwise spiralling, hyphal segments in filaments extend at a prescribed acute angle *θ_p_*, measured anticlockwise from the orientation of the major axis of the segment. This angle is known to be acute [22], but has not been previously quantified, and a key contribution of our work will be to infer a posterior distribution for this angle. The model features are summarised in Figure 4.

**Figure 3:**
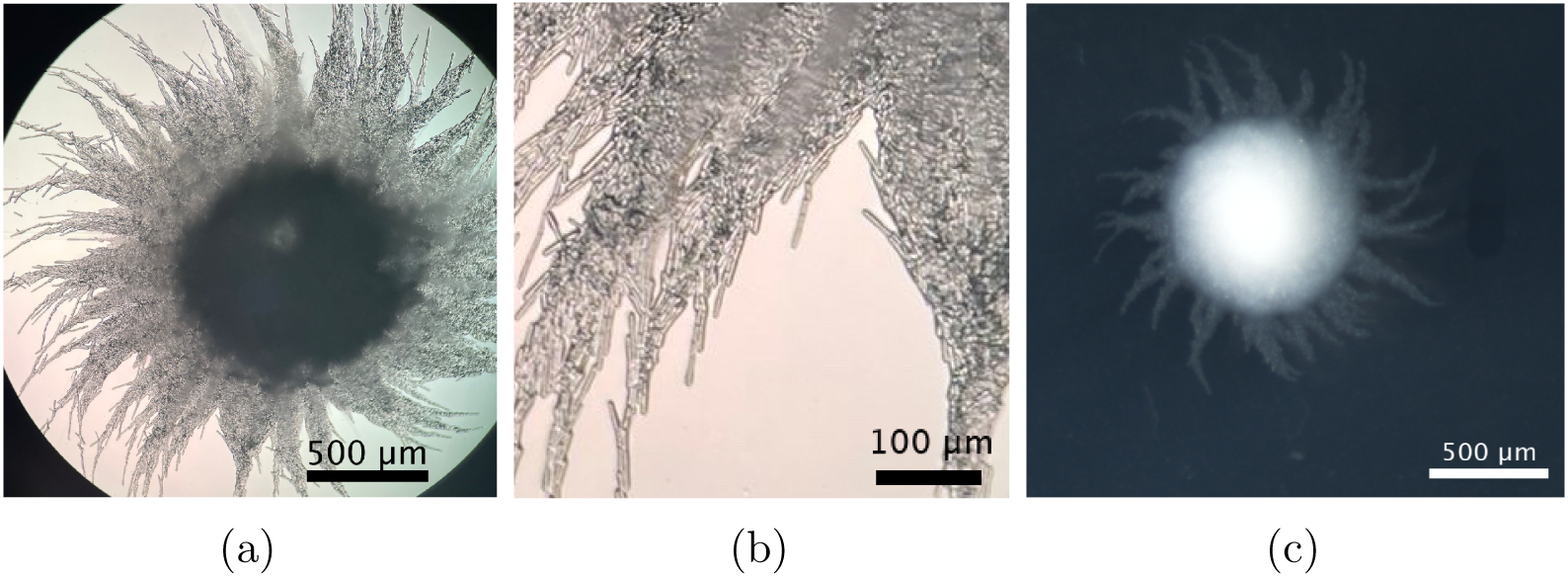
Images of *M. magnusii* at different magnifications, revealing the microscopic features of the colony structure when grown on CDS medium for seven days. (a) A colony exhibiting the spiral morphology. (b) Zoomed view of an experimental colony showing hyphae and conidia. (c) Full view of a *M. magnusii* colony after seven days of growth on YNB with high colony density. This colony is a different experiment to Figures 2, 3a and 3b, and is the colony used when inferring parameters to compare the model and experiment.

**Figure 4:**
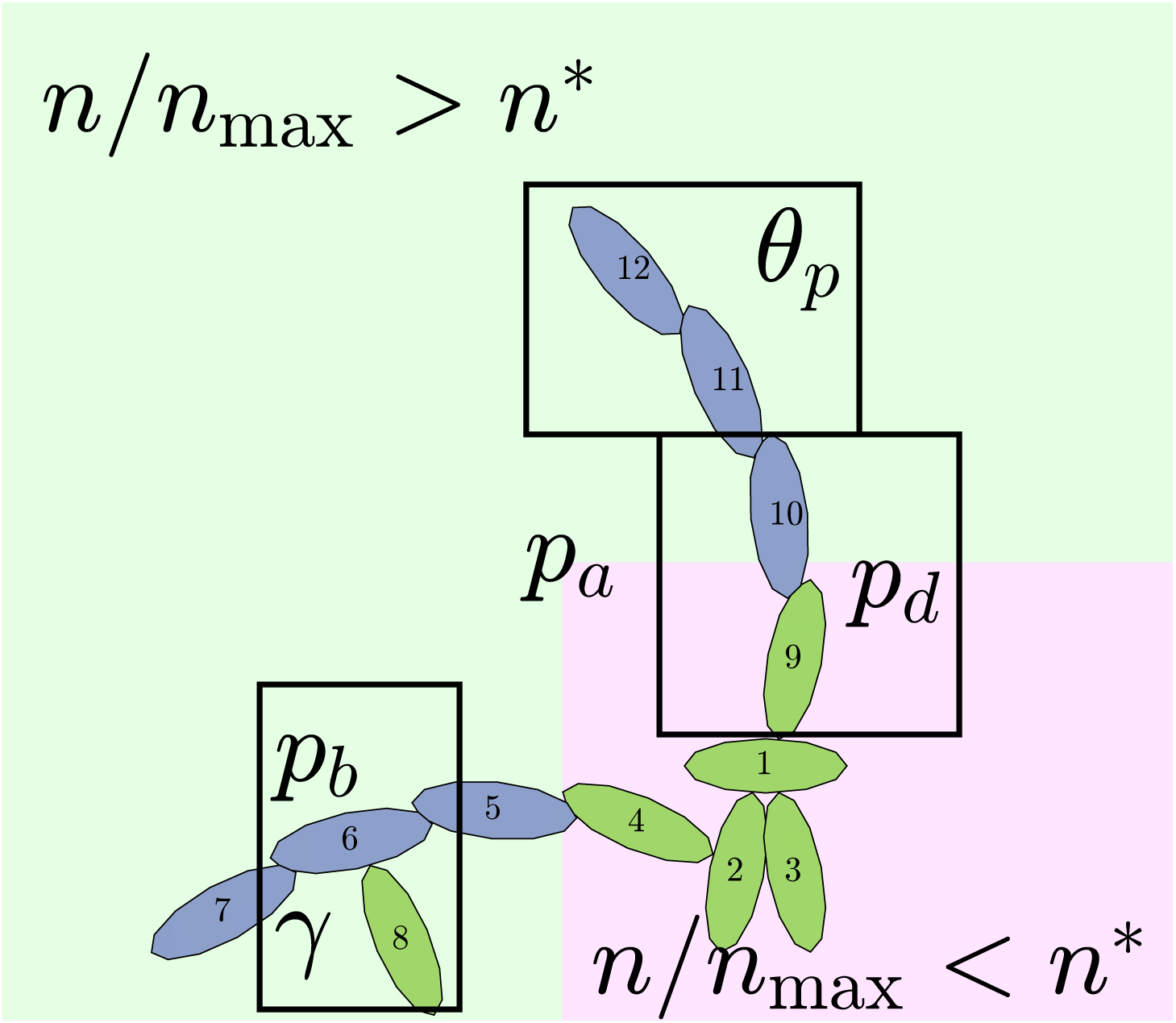
Cellular rules in the modified agent-based model. Regular hyphae that can extend from their sides are shown in green. Filament-forming hyphae that only extend at the angle *θ_p_* from their distal ends are shown in dark blue. The numbers represent a possible sequence of hyphal growth, starting from hyphal segment 1, that could generate this pattern.

We simulate hyphal growth one agent at a time until the colony reaches a prescribed number of agents, *n*_max_, chosen in advance such that the simulation ends when the simulated colony has approximately the same area as an experimental colony. Each step of the simulation involves selecting a hypha to produce a new segment, and proposing the type and location of the new agent. This behaviour depends on six parameters, ***θ*** = (*n*^∗^*, p_b_, p_d_, γ, p_a_, θ_p_*). After proposing the new agent, we implement partial volume exclusion to prevent two agents from occupying the same space. If the centre of a proposed new agent would lie within the boundary of another agent, the hyphal extension event does not occur.

On the length scale of *M. magnusii* experiments, the nutrient concentration is spatially uniform [68]. In the model, we use the existing colony area as a proxy for nutrient availability, such that hyphal filaments can emerge when the colony is sufficiently large. Consequently, we model neither time nor the nutrient concentration explicitly. The parameter *n*^∗^ ∈ [0, 1] is the threshold proportion of hyphae, relative to *n*_max_, which governs whether filament-forming hyphae can emerge. If the current number of agents in the simulated colony, *n*, is such that *n < n*^∗^*n*_max_, then only regular hyphae that can extend from both ends (shown in green in Figure 4) can be produced. When *n > n*^∗^*n*_max_, then filament-forming hyphae that extend from one end only (shown in dark blue in Figure 4) can emerge. The threshold *n*^∗^ thus represents the colony size, relative to *n*_max_, when the environmental conditions (*e.g.* low nutrient availability) required to produce hyphal filaments arise.

Once *n/n*_max_ *> n*^∗^, the other probabilities *p_a_, p_b_, p_d_*, and *γ* determine the next hypha to extend, and the type of new agent that is produced. The next agent selected to extend is governed by the probability *p_a_*. A regular hypha is selected with probability *p_a_*, and a filament-forming hypha is selected with probability 1 − *p_a_.* The particular agent to extend is chosen at random from all agents of the relevant type. This assumption represents the hypothesis that initiating filament formation is a colony-level decision.

If a regular hypha is selected to extend, then the resulting agent is filament-forming with probability *p_d_*, or regular with probability 1 − *p_d_.* If a filament-forming hypha is selected, then we first check whether that filament-forming hypha has an existing filament-forming daughter agent. If not, a new filament-forming agent is proposed. If there is an existing filament-forming daughter agent, then the behaviour is governed by *p_b_* and *γ.* First, with probability *γ* a different filament-forming agent without an existing filament-forming daughter is chosen to extend instead. Larger values of *γ* favour extended chains of individual filament-forming agents, and limit the appearance of regular hyphae within these chains. Then, if the agent with the existing filament-forming daughter proceeds to extend (which occurs with probability 1 − *γ*), the proposed new agent will be a regular hyphal segment with probability *p_b_*, or a filament-forming segment with probability 1 − *p_b_.* In the latter scenario, the proliferation event is aborted due to volume exclusion, because filament-forming agents can only extend at one site with orientation *θ_p_*. The flowchart in Figure 5 summarises steps taken to propose a new agent.

**Figure 5:**
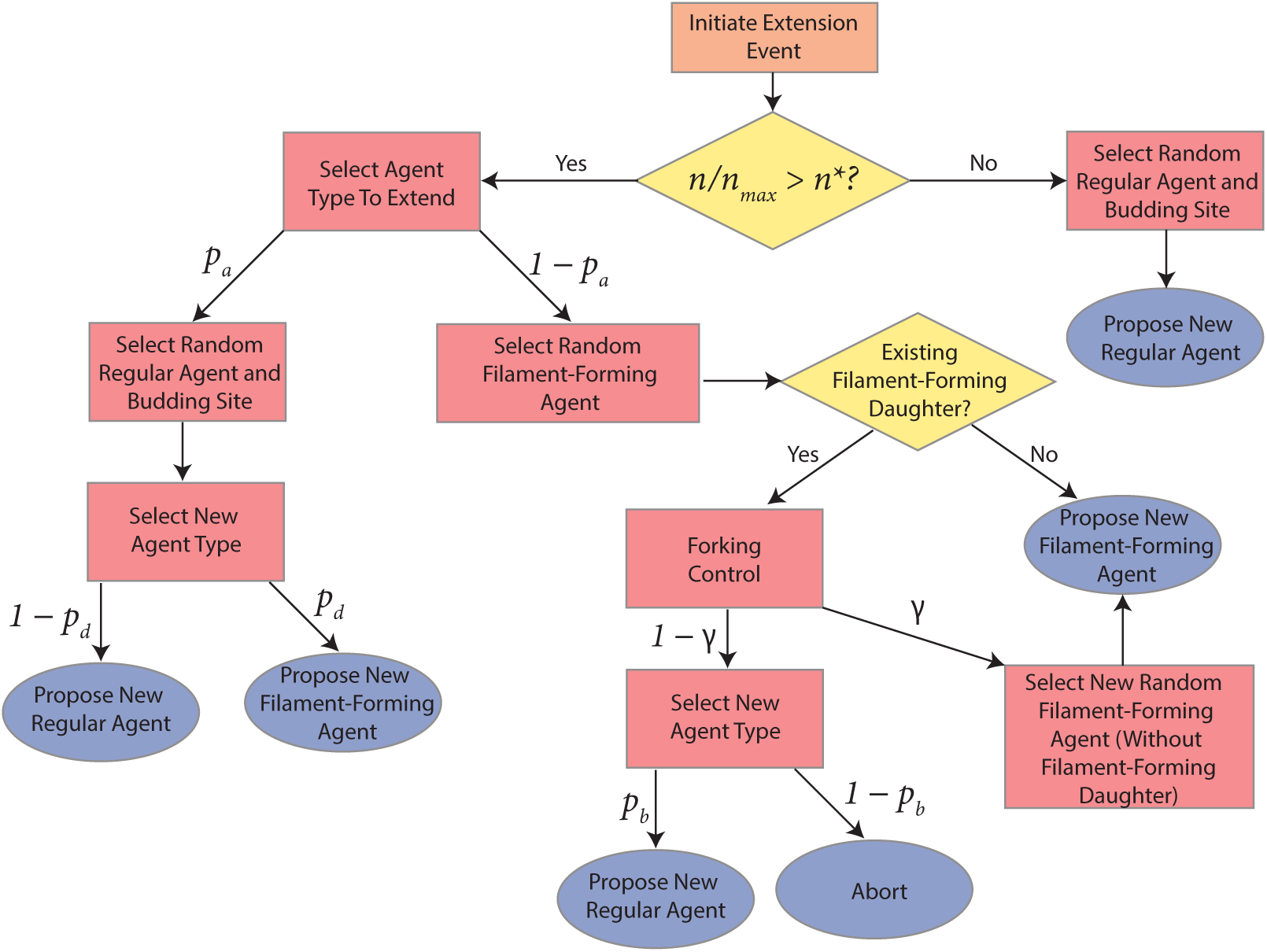
Flowchart describing the steps taken to propose a new agent (hyphal segment) in the agent-based model.

Since our agent-based modelling framework is similar to that applied to pseudohyphal growth [38], it is adaptable to other yeasts with different forms of growth. Should another yeast species that forms irregularly-shaped small colonies be studied, we could readily adapt this model, provided the agent size, budding patterns, and experimental colony area (in physical units) are known or assumed in advance. In Li et al. [38], we have already applied a similar model to multiple strains of *S. cerevisiae* grown in different environments. The *M. magnusii* model presented here is one of many possible model variations, and demonstrates how the model can also apply to hyphal growth.

### 3.2 Image processing and summary statistics

We process experimental photographs and simulation results to quantify the spatial patterns. We convert experimental photographs to binary images using the Tool for Analysis of the Morphology of Microbial Colonies (TAMMiCol) [66], such that black pixels indicate regions occupied by the colony, and white pixels indicate unoccupied regions. The binary image for the *M. magnusii* colony is in Figure 6b. To enable direct comparison, we save images of the simulated colonies at the same scale and resolution as experimental photographs. We simulate colonies until they attain a target number of agents, which we prescribe. To obtain the target number of agents for the simulations that best fits the experimental data, we used the golden section search to minimise the difference in colony area between the simulated and experimental colonies.

**Figure 6:**
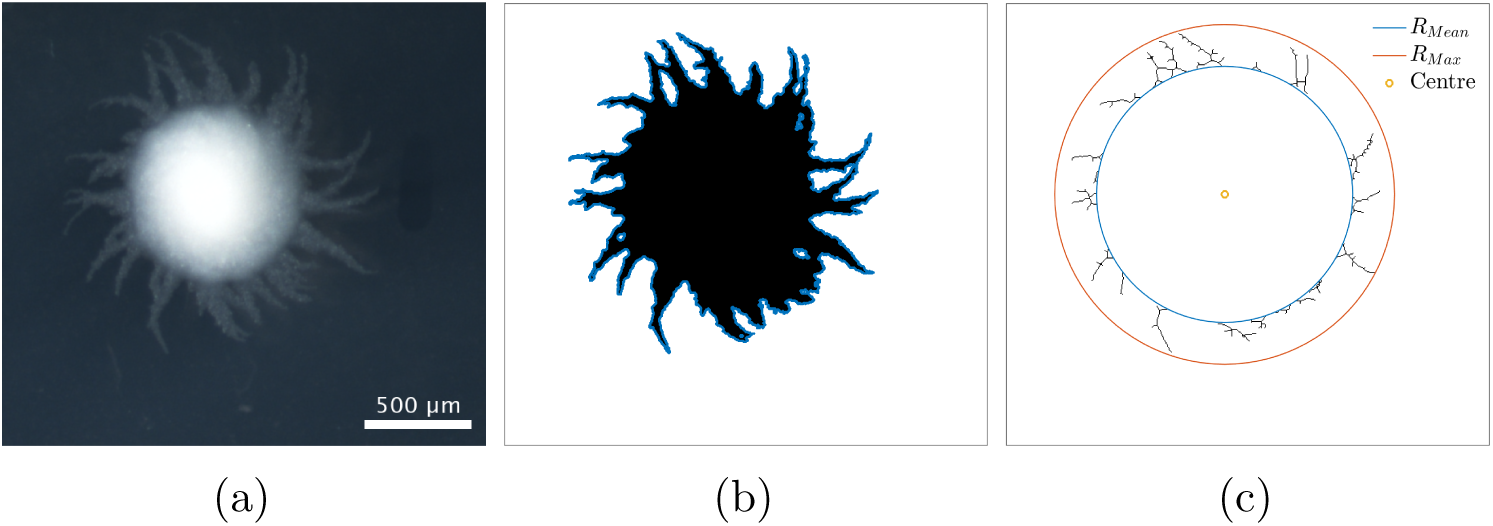
Visualisation of the summary statistics used in this paper. (a) Original experimental photograph of a *M. magnusii* colony with spiral hyphae. (b) Binary image with area overlayed with perimeter (blue). (c) Mean and maximum radii overlayed with the skeletonisation of the binary image in (b).

For both experimental and simulation results, we define five spatial statistics to characterise morphology: maximum radius (*R*_max_), mean radius (*R*_mean_), filamentous area 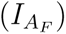, branch count (*I_B_*), and compactness (*I_C_*). The two radii provide two measures of colony size. The maximum radius is the largest distance from the colony centroid to an occupied pixel, and the mean radius is the average distance between the centroid and all pixels on the colony perimeter. The colony centroid is found using Matlab’s **regionprops()** function. Filamentous area is the number of occupied pixels between the mean and maximum radii, and therefore quantifies the extent of the non-uniform hyphal growth. Branch count quantifies the complexity of the hyphal filament pattern, and is the number of segments identified in a skeletonised image produced using Matlab’s `bwmorph()` function, that lie between the mean and maximum radii. Figure 6c provides an example of a skeletonised image. Compactness is the ratio between the colony perimeter and the perimeter of a circle with the same area as the colony, and quantifies the shape irregularity of the colony. Compactness is defined as *I_c_* = *P/*(4*πA*), where *P* is the number of occupied pixels on the colony perimeter (including holes within the colony), and *A* is the total number of occupied pixels in the colony. The perimeter is determined using Matlab’s `bwperim()` function. To ensure that each summary statistic has approximately equal weighting in the parameter inference, we ensure that each statistic only takes values on the unit interval by dividing each summary statistic by the maximum observed value of that statistic from 2000 simulations of the agent-based model.

### 3.3 Inference using sequential neural likelihood estimation

We do not have a high-resolution photograph of the entire colony that we analyse. Consequently, our model contains parameters governing an agent’s behaviour that cannot be observed on the microscopic scale of a single agent. Therefore, we need to infer these parameters by comparing the macroscopic output of the model with experiments. Due to the high-dimensional output of our model, we do not have closed-form expressions for the likelihood function. So, we use likelihood-free Bayesian inference to compute approximate posterior distributions for the model parameters given an experimental photograph. We adopt a sequential neural likelihood estimation (SNLE) method [51, 3]. This method uses simulated data to train a conditional density estimator for the likelihood function. We can then sample from the posterior distribution using a standard Metropolis–Hastings Markov Chain Monte Carlo (MHMCMC) approach [27], by substituting the approximate likelihood for the true function. The sequential naming of the algorithm refers to how the likelihood approximation is trained over a number of rounds, such that at the end the majority of the data is from areas of high posterior density resulting in an accurate approximation [51].

We use a Gaussian mixture density network (MDN) as our surrogate likelihood model [10]. The parameters ***θ*** = (*n*^∗^*, p_b_, p_d_, γ, p_a_, θ_p_*), are constrained to [1, 0]^5^ × [0*, π/*20], hence we apply a logit transform and fit the surrogate likelihood model in an unconstrained space. Each round of training used 1,000 simulations and then the sampling from the resulting posteriors was performed using an MH sampler. The initial training data was sampled uniformly from [1, 0]^5^ × [0*, π/*20] and the priors were taken to be the same. Four rounds of training were required to achieve sufficient convergence of the approximate posterior. More details on the training procedure are provided in the Appendix. See also Li et al. [37] for a similar procedure, but using only a single round of training.

## 4 Model simulation and parameter inference results

We perform the inference on a single experimental image to avoid issues of inter-experiment variability. A pair plot of the posterior distribution is shown in Figure 7. Figure 8 shows prior and posterior predictive distributions for the five summary statistics, and Figure 9 shows simulations from eight sets of parameters randomly sampled from the posterior. The observed summary statistics are in all cases in the tails of the prior predictive distribution, but the posterior predictive distributions’ modes are mostly centred on them, indicating that the inference algorithm is performing correctly. We also present the posterior from the initial round of training in the Appendix to demonstrate that the posterior from the initial round is much wider and less informative than the posterior after 4 rounds. Figure 9 provides an additional visual demonstration that the inferred parameters are capturing the correct morphology, with patterns resembling the experimental image.

**Figure 7:**
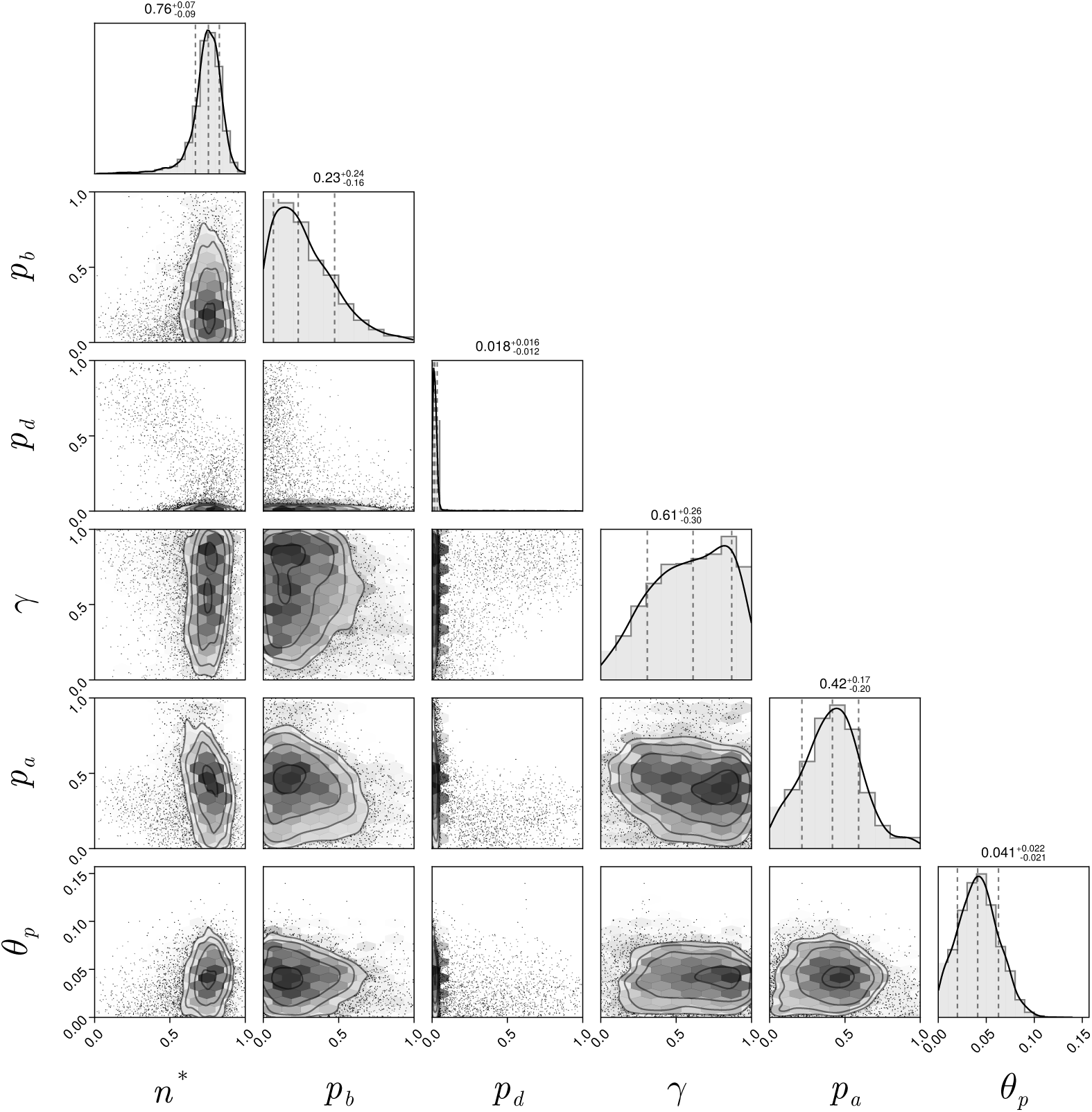
Pair plot of posterior distributions after three iterations of the sequential neural likelihood estimation. All priors were uniform over the axes ranges shown in the figure.

**Figure 8:**
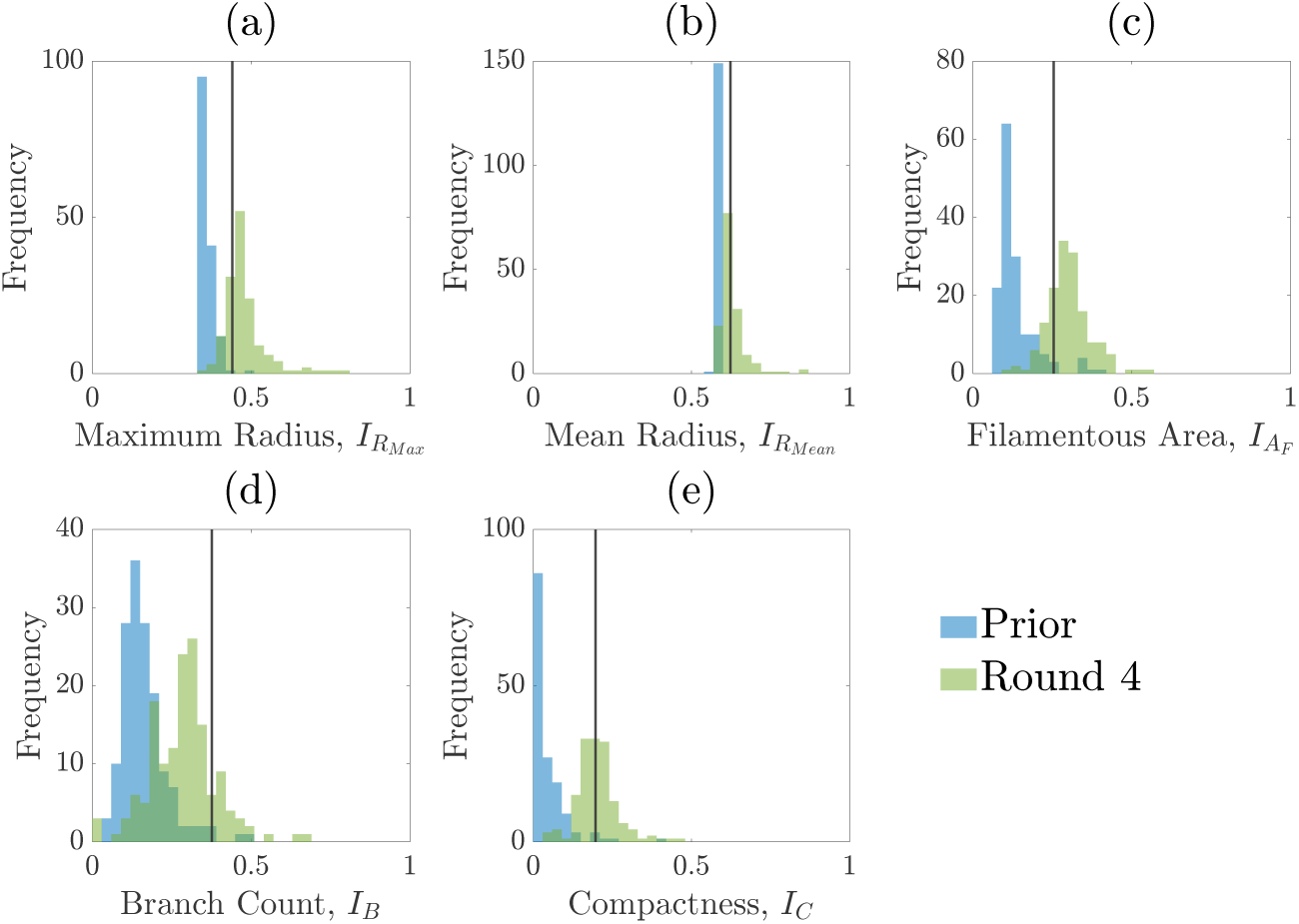
Normalised summary statistics generated from sampling from either the prior or from the posterior distribution after four rounds of SNLE. The vertical line represents the summary statistic from the experimental colony shown in Figure 6. Blue histograms are of the summary statistics for simulations sampled from the prior, and the green histograms are for simulations sampled from the posterior distribution after round 4 of the SNLE. (a) Maximum radius (b) Mean radius (c) Filamentous area (d) Branch count (e) Compactness.

**Figure 9:**
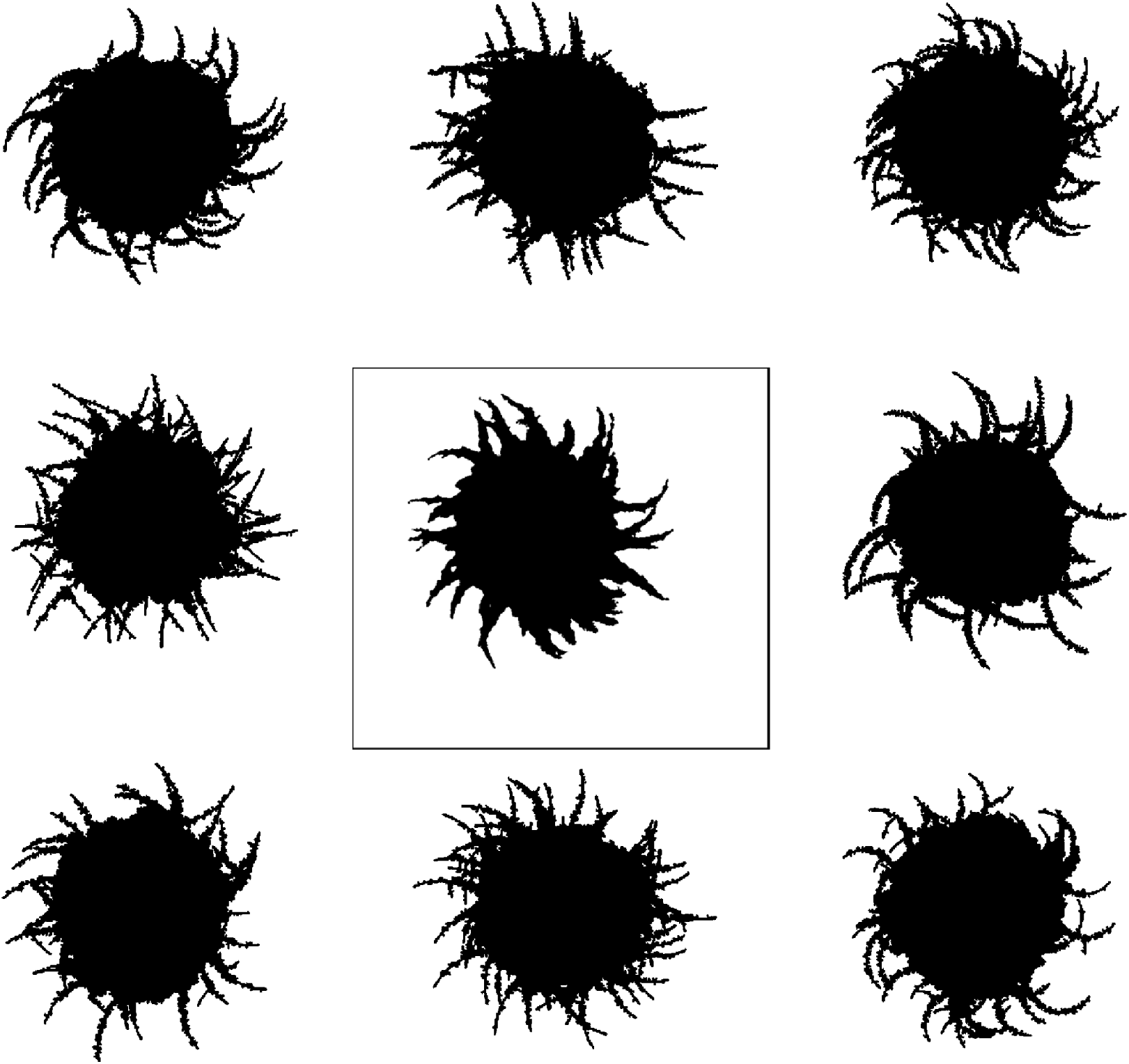
A binary image of an experimentally-produced colony of *M. magnusii* grown on YNB containing 150 mm ammonium sulfate (centre panel), flanked by eight simulations drawn from the posterior from Figure 7.

The data are informative for all parameters except *γ*. The parameters *n*^∗^ and *p_d_* are strongly identified, whereas the marginal posteriors for *p_b_, p_a_*, and *θ_p_* are broad, meaning that these parameters are only weakly identified. As expected, the threshold for initiation of spiral filamentation, *n*^∗^, is inferred well with a tight marginal posterior with mode of 0.76. This parameter is essential in controlling when filamentation begins, which is important in determining the overall morphology and size of the colony. The parameter *p_d_*, which is the probability that a filament-forming agent emanates from a regular agent, is strongly inferred to be small. If *p_d_* is small, we expect fewer longer spiralling filaments, whereas a large *p_d_* would produce more, shorter spiralling filaments. From the experimental images, we anticipate smaller values of *p_d_* because filaments are distinct and long and this is confirmed by the marginal posterior that puts almost all mass on values *p_d_ <* 0.1.

Another crucial parameter in our model is the angle between successive segments, *θ_p_*, as this determines the amount of spiralling. This parameter could be identified, and its mode is 0.041 radians (2.3 degrees). This result provides an estimate of the angle, which has not previously been quantified for *M. magnusii*. Although the angle is small, it still gives rise to observable colony-scale spiral patterns, as Figure 9 shows. Since *θ_p_* is constant in the model, within each simulated colony the spirals tend to have similar arc radii, with some variation that reflects the variation in the experimental image. The model can also reproduce characteristic microscopic patterns within the filamentous regions themselves (see Section A in the Appendix for further details).

The probability *p_a_* governs whether a regular or filament-forming hypha is chosen to produce a new segment. This parameter is weakly identified, suggesting that capturing the initial emergence of filament-forming hyphae and the angle between successive segments are more important for predicting the morphological features captured by the summary statistics. Similarly, *p_b_*is weakly identified. Changes to this parameter influence the thickness of the filaments by allowing regular hyphae to form alongside filament-forming ones. Since our summary statistics do not contain a metric that explicitly quantifies the width or curvature of filaments, this parameter cannot be well inferred. The data are uninformative for the parameter *γ.* This parameter, which controls forking from filament-forming segments, was important for the pseudohyphal *S. cerevisiae* yeast growth in Li et al. [38], but is poorly identified for these hyphal *M. magnusii* colonies. These results suggest that the value of *γ*, and hence the forking control process, is less important for determining the *M. magnusii* morphology.

The statistics for maximum radius, mean radius, and compactness are especially well-predicted by our posterior predictive simulations (see Figure 8). There is larger variability in the filamentous area and branch count among the simulations, although overall the simulations still align well with the experiment. Variability in both the filamentous area and branch count is consistent with the finding that parameters affecting filament width are only weakly identified due to the summary statistics not capturing filament width.

## 5 Other *M. magnusii* yeast morphologies

In different experimental conditions, *M. magnusii* colonies form different morphologies. We explore these morphologies through additional simulations of the model. In a nutrient-rich environment, the yeast tends to grow in a rounded hexagonal shape as shown in Figure 10a, reminiscent of Edenlike growth [24]. Although the round colony does not exhibit irregular filamentation, the almost hexagonal perimeter we see in the simulation mimics the experiment, which could be due to the shape and pattern of growth for regular hyphae. Under other conditions, *M. magnusii* has also been shown to have straight filaments of varying thicknesses as shown in Figures 10b and 10c. These patterns are more similar to the patterns formed by hyphal colonies of the important pathogenic yeast *C. albicans* [8, 19, 47]. With specifically-chosen parameter values, we can also capture these patterns in model simulations, as presented in Figures 10d–10f.

**Figure 10:**
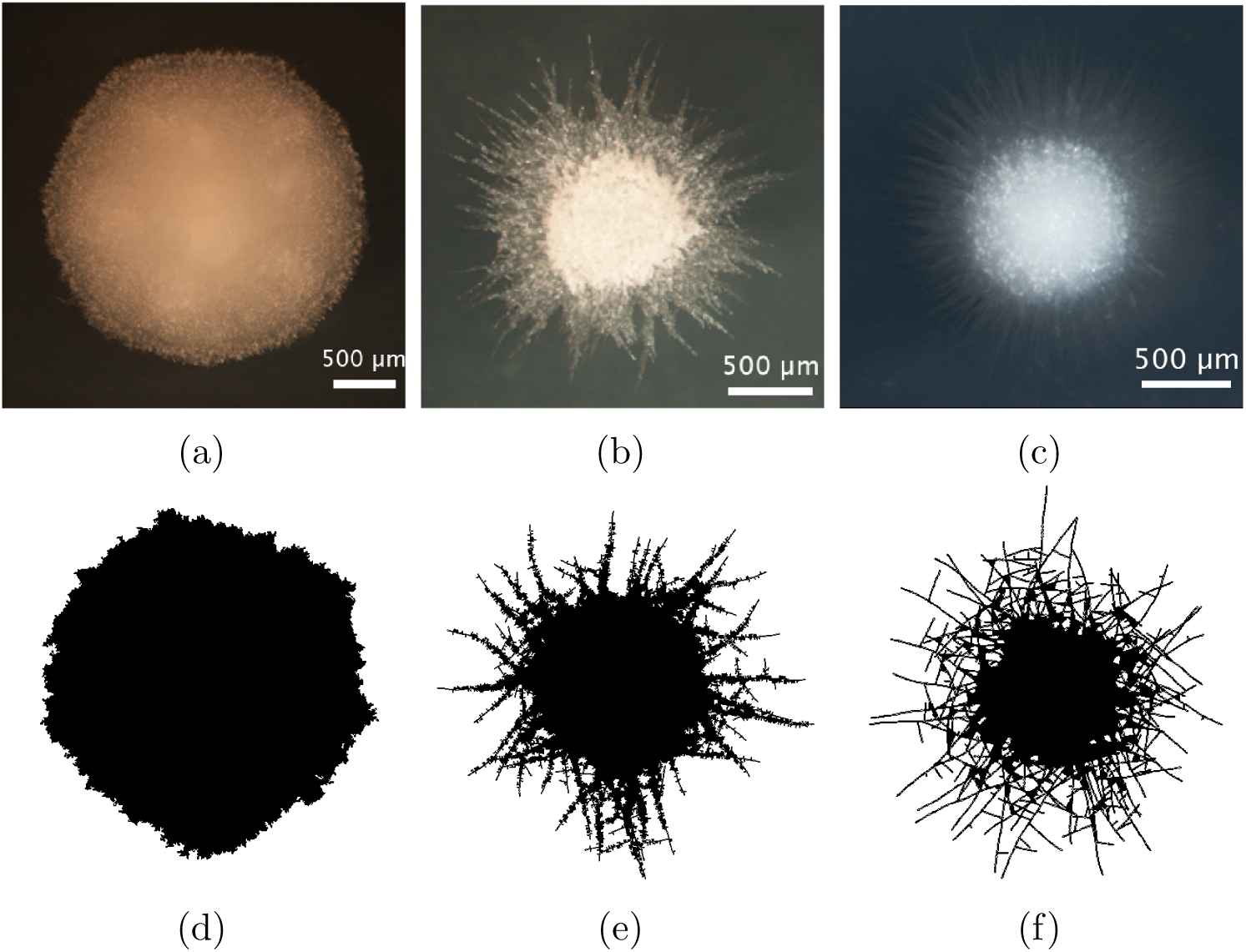
Other *M. magnusii* yeast morphologies under varying environmental conditions. (a) Experimental colony grown on CDS with low yeast density. (b) Experimental colony grown on CDS at high yeast density. (c) Experimental colony grown on YNB at high yeast density. (d) Simulation with *n*_max_ = 10, 000 agents and parameters *n*^∗^ = 1, *p_a_* = 0.25, *p_b_* = 0.3, *p_d_* = 0.1, *γ* = 0.05, and *θ_p_* = 0.05. (e) Simulation with *n*_max_ = 10, 000 agents and *n*^∗^ = 0.6, *p_a_* = 0.25, *p_b_* = 0.3, *p_d_* = 0.1, *γ* = 0.05, and *θ_p_* = 0.05. (f) Simulation with *n*_max_ = 10, 000 agents and *n*^∗^ = 0.5, *p_a_* = 0.1, *p_b_* = 0.02, *p_d_* = 1, *γ* = 0.90, and *θ_p_* = 0.05.

The ability to capture multiple experimentally-feasible *M. magnusii* morphologies suggests that our off-lattice model is consistent with the biology of hyphal growth. Furthermore, differences in parameter values across the three simulations in Figure 10 may indicate how the environment influences *M. magnusii* growth. In the hexagonal colony of Figures 10a and 10d, the threshold for filament-forming hyphae is set to *n*^∗^ = 1, whereas this parameter is *n*^∗^ = 0.6 and *n*^∗^ = 0.5 for the other colonies. The experimental colony in Figure 10a was grown on CDS media with low density of colonies within the culture well, whereas the colonies in Figures 10b and 10c were grown under conditions of high colony density. The differences in *n*^∗^ indicate that *M. magnusii* does not experience low-nutrient stress in the low colony density case, but does in high-density environments, where more colonies compete for the same nutrients. These results are consistent with the hypothesis that the transition between regular and filament-forming hyphae is mediated by nutrient availability. In contrast, the proliferation angle *θ_p_*is held constant across all three simulations in Figures 10d–10f. Since *θ_p_*was identifiable in the parameter inference, this angle could be an intrinsic feature of *M. magnusii* growth regardless of environment. Finally, although *p_b_* and *γ* were difficult to identify, comparing the simulations in Figures 10e and 10f reveals their impact. The simulation with larger *p_b_*, smaller *γ*, and smaller *p_d_*favours thicker hyphal filaments (Figure 10e), because regular hyphal segments are more likely to occur within filaments. Conversely, the simulation with smaller *p_b_*, larger *γ*, and larger *p_d_* is more likely to produce filaments, but less likely to contain regular hyphae within filaments, creating the thinner filaments seen in Figure 10f.

## 6 Discussion and conclusion

In this paper, we have presented an off-lattice agent-based modelling and parameter inference procedure that yields biological insight into small yeast colonies formed in harsh environments. We applied this procedure to a new strain of *M. magnusii* recently bioprospected from a Tasmanian cider gum tree. Since this strain produced a spiral-like morphology, we adapted our agent-based model to incorporate hyphal segments and extension at a constant angle between successive segments. We used image processing and spatial statistics to compare the shape of simulated colonies with the experiment. Using Bayesian inference, we estimated the mean angle between segments to be 2.3^◦^ [1.1^◦^, 3.6^◦^] (95% credible interval), a result that has not previously been quantified. One potential limitation of our method is that the termination criterion was tuned to match the observed area while size-based summary statistics were also used, which might produce overconfident posteriors. However, colonies simulated using the inferred parameters closely resembled those from the experiment. We also extended this result further to obtain simulations of other experimentally-realised shapes with different parameters. We also provide open-source code for the agent-based models and parameter estimation techniques on GitHub.

Our work provides a framework for understanding the colony patterns formed by new yeast strains uncovered by bioprospecting. Adapting the agent-based model to meet the biological assumptions and performing parameter inference will allow researchers to gain insight into the key cell-level mechanisms underpinning the patterns. Improvements for future work could also include a summary statistic to capture the thickness of branches, which might make the parameter *p_b_* more informative. Additionally, this paper has reviewed several forms of surface growth in yeast colonies. However, in some experiments yeast cells grow on the surface and also penetrate into the agar medium [20]. This invasive growth is of great interest to biologists due to its importance for pathogenic infections. In future work, we intend to extend the modelling and parameter inference methods to three-dimensions to investigate this invasive growth phenomenon. From the biological perspective, the environmental and genetic drivers of spiralling in *M. magnusii* remain to be fully understood. In the mould fungus *Neurospora crassa*, the gene *coil-1* promotes spiralling or coiling [7], and yeast morphologies broadly depend on genetic and environmental factors [31]. Future experimental work could involve disrupting specific genes or growing *M. magnusii* in varying conditions, to assess how these factors impact spiral formation.

## Acknowledgements

KL acknowledges funding from the Australian Government through a Research Training Program (RTP) Scholarship. VJ, JEFG, BJB, and AKYT acknowledge funding from the Australian Research Council (Grant numbers DP230100406, and DE240100097). The authors acknowledge supercomputing resources from The University of Adelaide’s maths1 High Performance Computing (HPC) service.

## Code availability

All code and data are available on GitHub at: https://github.com/kaili2019/Li2025-Spiral-Yeast.

## A Zoomed simulation result

Figure 11 compares the microscopic detail in the filamentous region between an experiment and a simulation of *M. magnusii*. These zoomed images indicate that the agent-based model can successfully represent the spiral filament-forming process.

**Figure 11:**
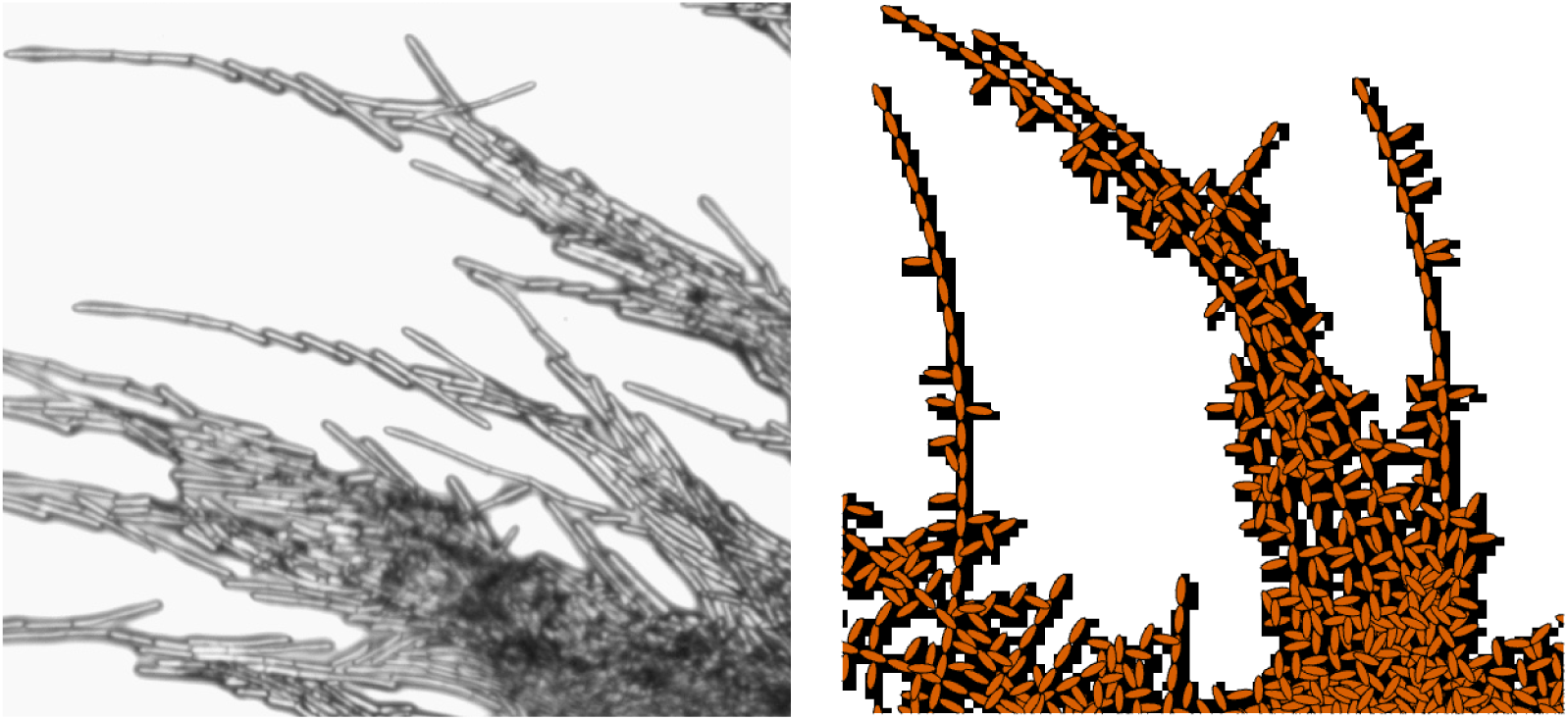
Comparison of an experiment and simulation (not to scale). Left: zoomed view of the filamentous region of a *M. magnusii* colony from one of our experiments. Right: A typical agent-based simulation, zoomed in near a hyphal filament. Brown ellipses indicate the locations of hyphal segments, and the black overlay indicates the simulated colony after binarisation.

## B Multiple agent sizes

We also investigate the impact of agent shape on the spiral yeast morphology. To demonstrate the colony morphology is robust to changes in agent shape, we simulate colonies where the regular hyphal segments and filamentous hyphal segments have different geometry. We use the same parameter values as in Figure 9, varying only the agent shape. There is minimal difference between the three colonies, suggesting that other parameters such as *θ_p_* drive the spiral colony morphology in *M. magnusii*.

**Figure 12:**
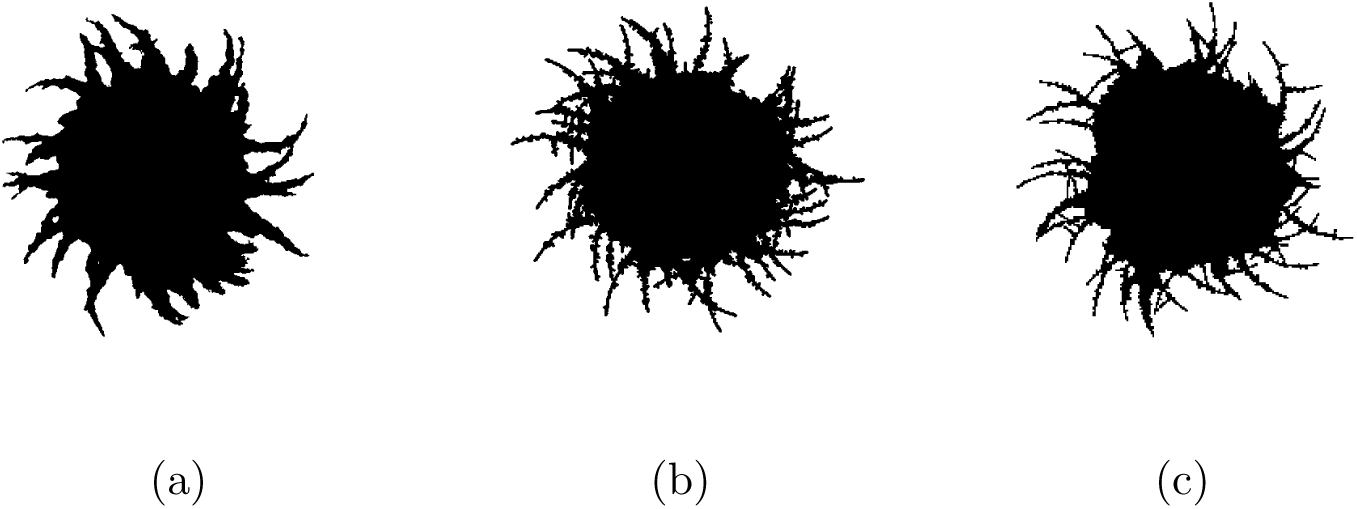
(a) Experiment of *M. magnusii* grown on YNB medium. (b) Spiral yeast simulation with parameters *n*^∗^ = 0.76, *p_a_* = 0.42, *p_b_* = 0.23, *p_d_* = 0.02, *γ* = 0.77, and *θ_p_*= 0.04, drawn from the posterior distribution. All agents have the same size, with aspect ratio 0.33. (c) Spiral yeast simulation with the same parameters as (b), but two possible agent sizes, with aspect ratios 0.75 (regular) and 0.33 (filament-forming). Each agent has the same area.

## C Additional parameter inference results

The following figures summarise the summary statistics obtained after each round of sequential neural likelihood applied to the experimental image.

**Figure 13:**
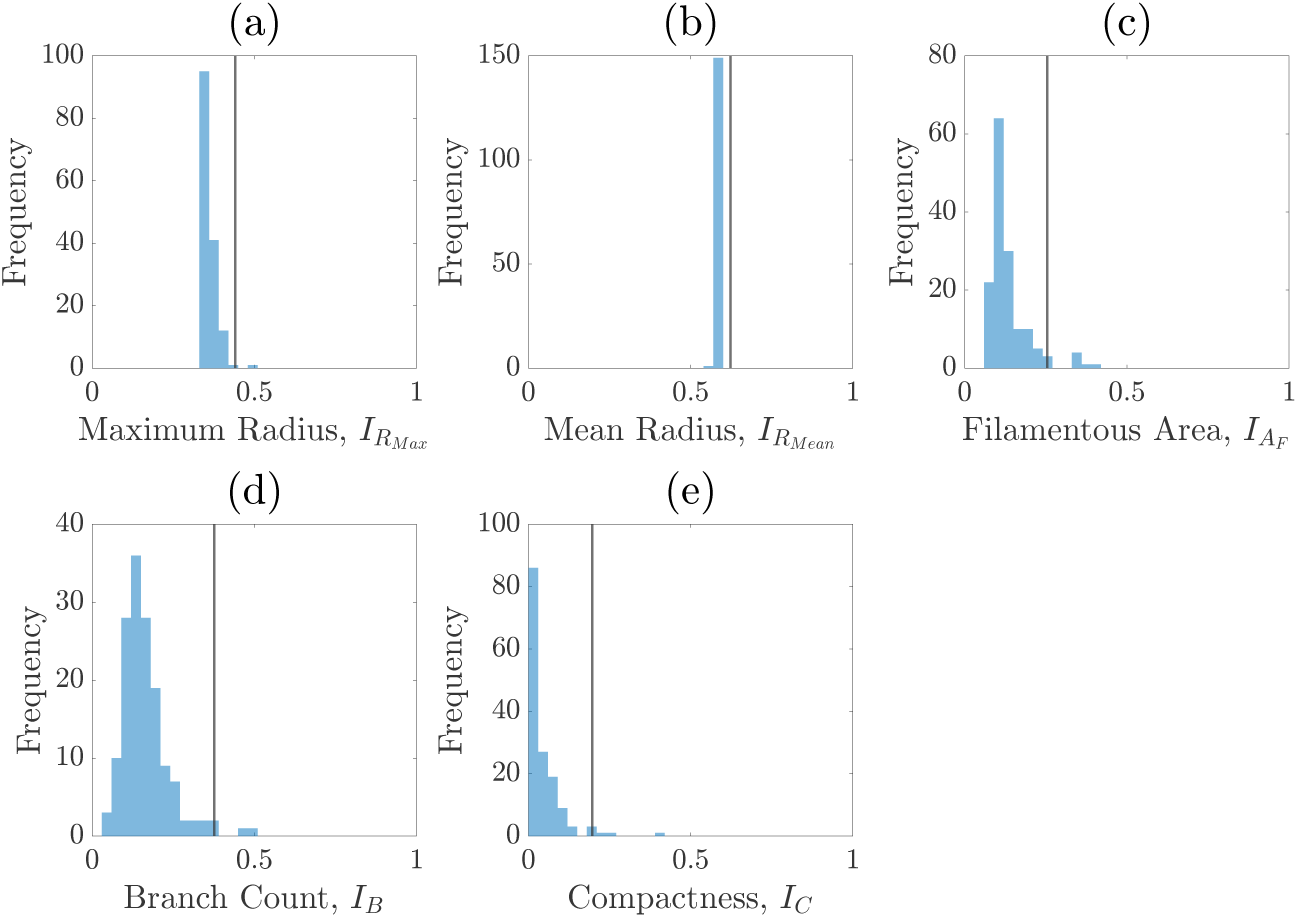
Summary statistics of the priors.

**Figure 14:**
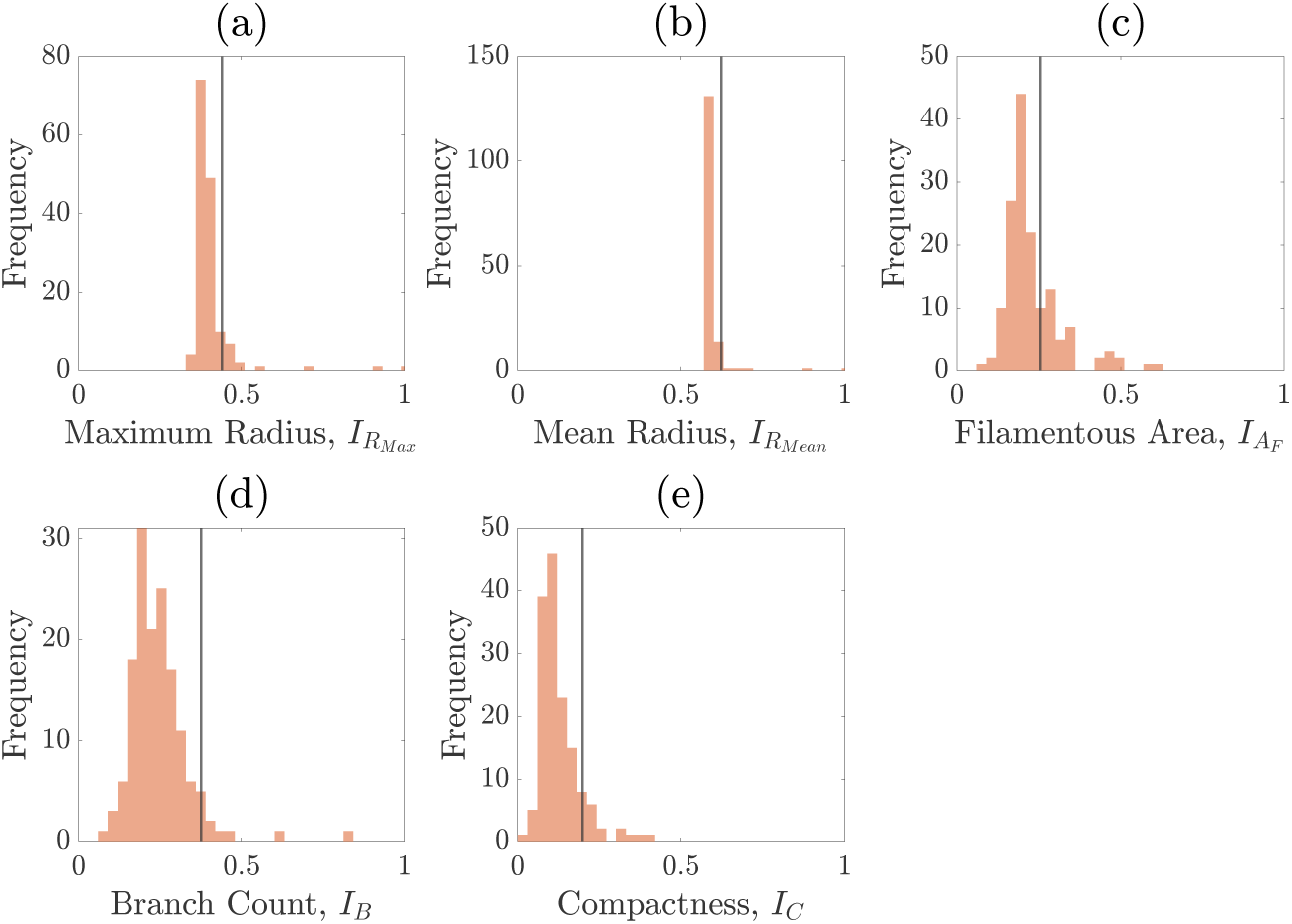
Summary statistics of round one.

**Figure 15:**
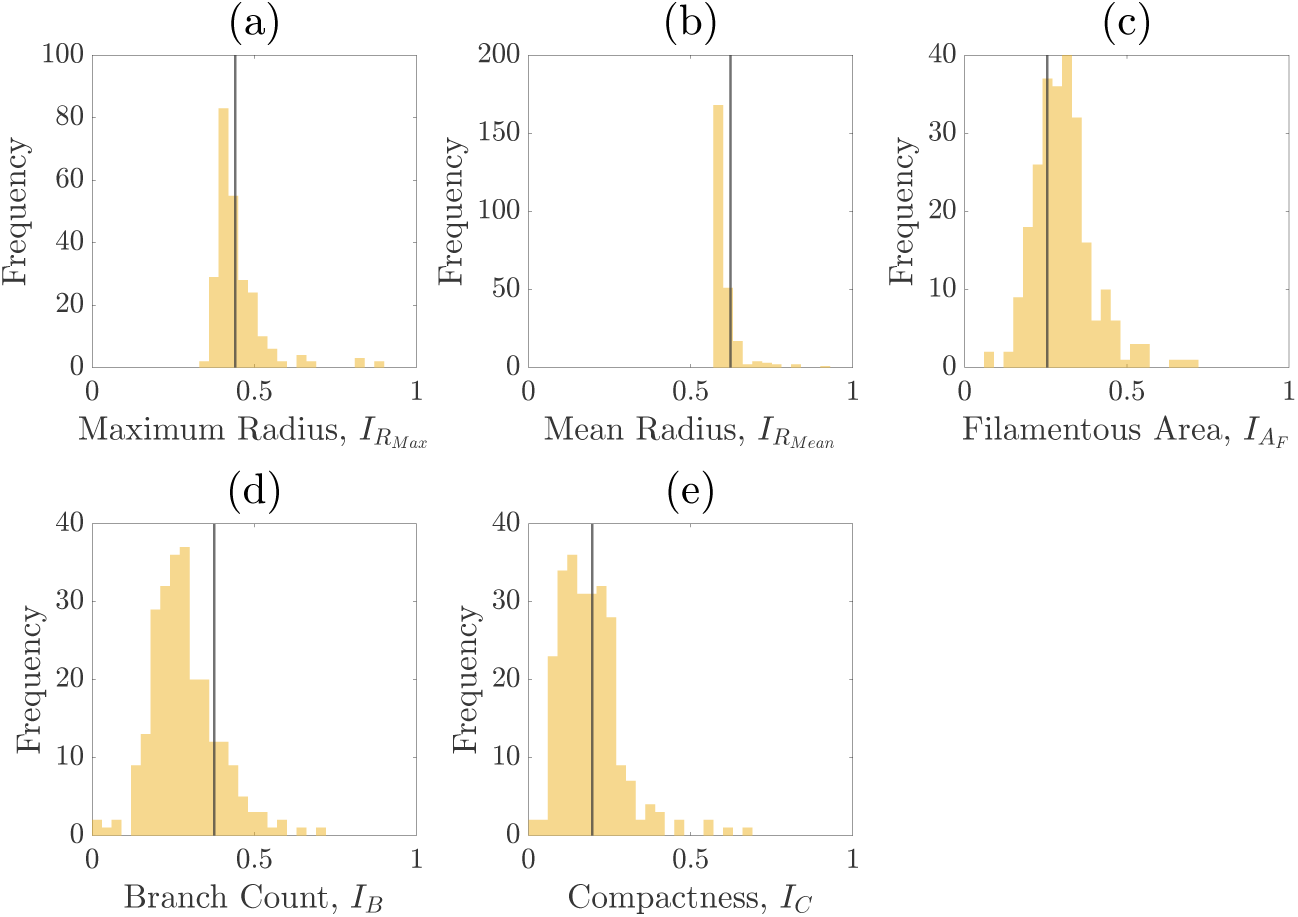
Summary statistics of round two.

**Figure 16:**
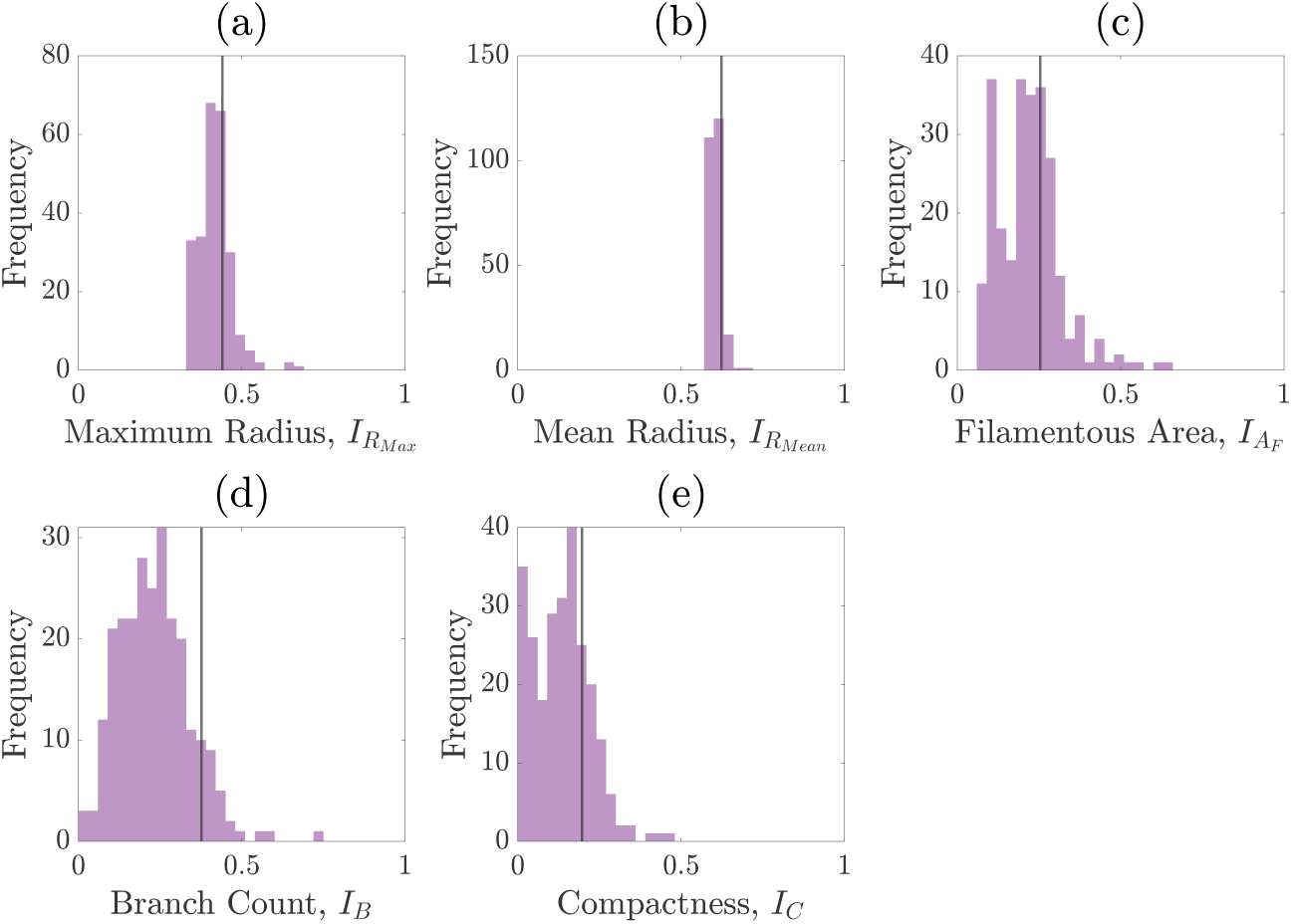
Summary statistics of round three.

**Figure 17:**
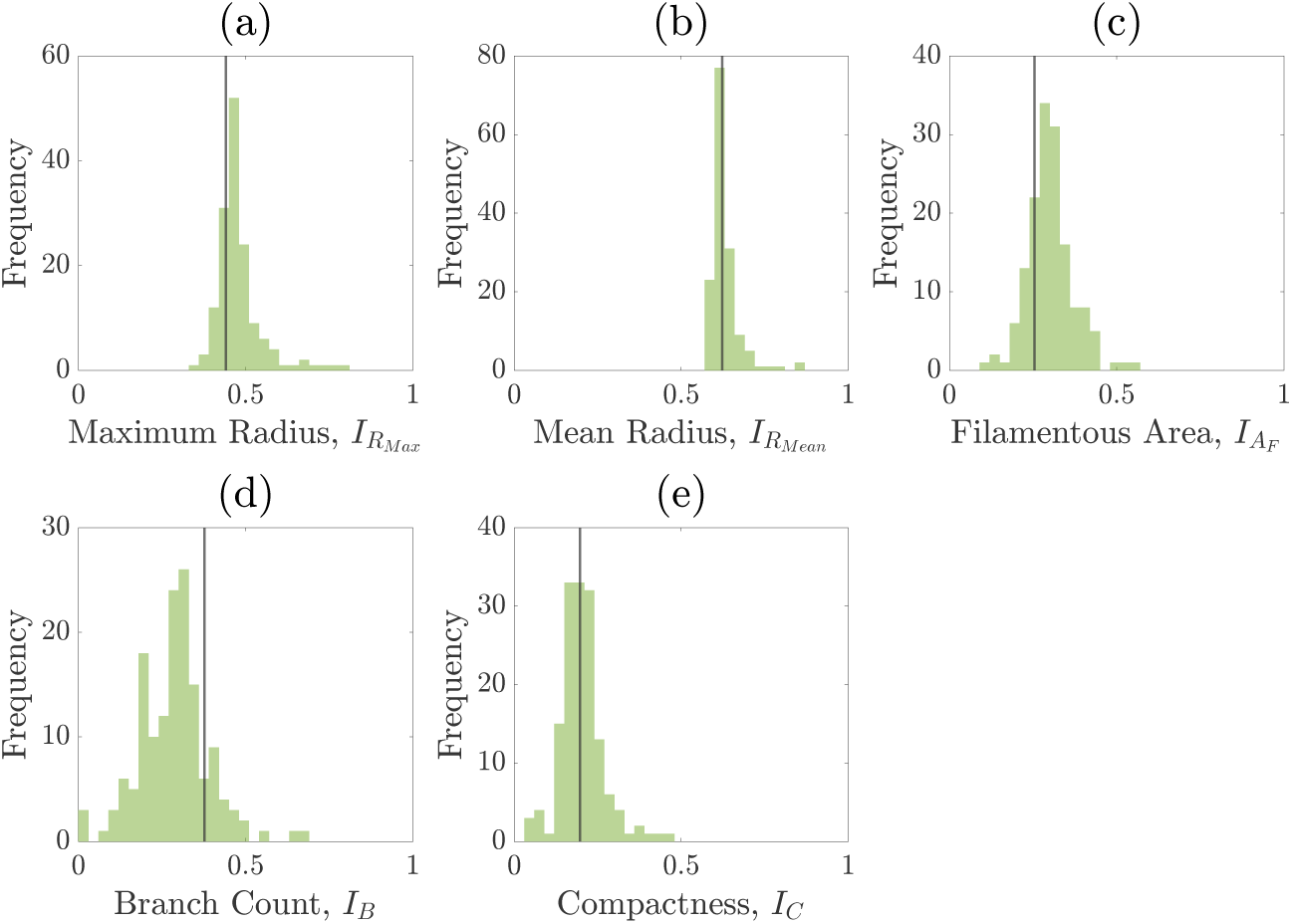
Summary statistics of round four.

**Figure 18:**
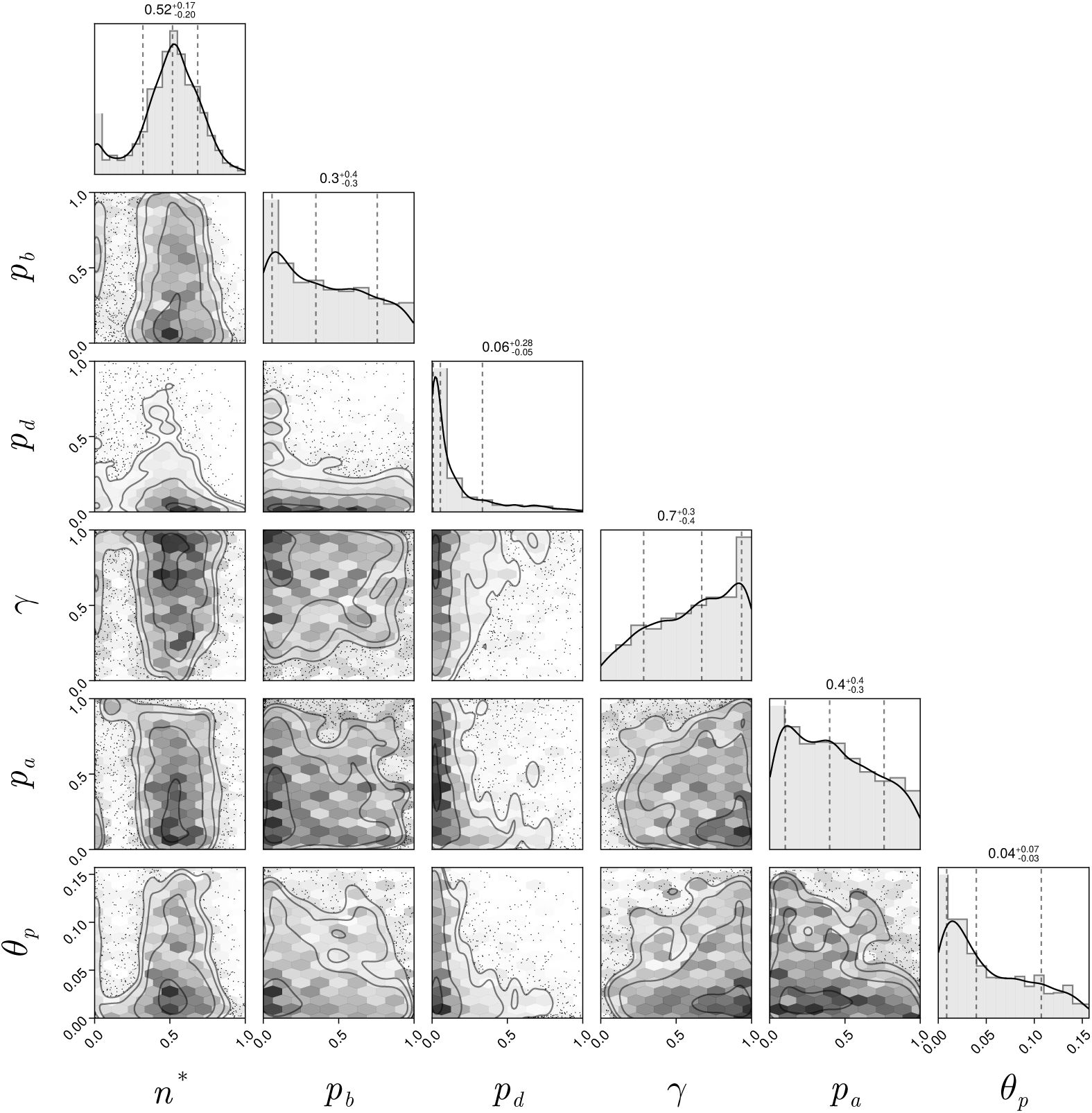
Pair plot of posterior distributions after initial round of the neural likelihood estimation.

## D SNLE details

We use a Gaussian mixture density network (MDN) as our surrogate likelihood model [10], which is implemented using the MixtureDensityNetworks.jl package in Julia. The parameters ***θ*** = (*n*^∗^*, p_b_, p_d_, γ, p_a_, θ_p_*), are constrained to [1, 0]^5^ × [0*, π/*20], hence we apply a logit transform and fit the surrogate likelihood model in an unconstrained space. Posterior sampling was also carried out in this space and then the resulting samples were transformed back to constrained space for visualisation and analysis.

The architecture of the MDN was chosen to be the same as in Li et al. [37], which worked well for a similar model and posterior geometry. The MDN had two hidden layers of width 64 and six mixtures. The MDN was trained using the Adam optimiser with a learning rate of 1 × 10^−3^, a batch size of 16 and for 600 epochs. Convergence was assessed by examining the loss over the number of epochs.

For each round of training a dataset of 1,000 simulations from the ABM was generated, which is equivalent to approximately 24 hours of high-computing time. Then, posterior sampling was performed using a basic Metropolis– Hastings algorithm. This used a basic Gaussian random walk proposal with standard deviation 1. We generated 200,000 MCMC samples to ensure adequate convergence of the posterior distribution, discarding the first 10,00 samples as burn-in and applying a thinning factor of 10 to reduce autocorrelation, resulting in an effective sample size exceeding 550 for each parameter. Owing to the simple geometry of the posterior in each round, the Markov chain mixed rapidly, requiring minimal manual tuning. It was found that 4 rounds of training were sufficient for convergence of the approximate posterior, which was assessed visually by changes in the distribution over rounds as well as the distributions of the summary statistics.

